# Neural oscillatory activity and connectivity in children who stutter during a non-speech motor task

**DOI:** 10.1101/2022.10.19.512866

**Authors:** Valeria C Caruso, Amanda Hampton Wray, Erica Lescht, Soo-Eun Chang

## Abstract

Neural motor control rests on the dynamic interaction of cortical and subcortical regions, which is reflected in the modulation of oscillatory activity and connectivity in multiple frequency bands. Motor control is thought to be compromised in developmental stuttering, particularly involving circuits in the left hemisphere that support speech, movement initiation and timing control. However, to date evidence comes from adult studies, with limited understanding about motor processes in childhood, closer to the onset of stuttering. In this study, we investigated the neural control of movement initiation in children who stutter and children who do not stutter by evaluating transient changes of EEG oscillatory activity and connectivity during a simple button press motor task. We found reduced modulation of left hemisphere oscillatory power, phase locking to button press and phase connectivity in children who stutter compared to children who do not stutter, consistent with previous findings of dysfunction within the left sensorimotor circuits. Interhemispheric connectivity was also weaker at lower frequencies (delta, theta) and stronger in the beta band in children who stutter than in children who do not stutter. Taken together, these findings indicate weaker engagement of the contralateral left motor network in children who stutter even during low-demand non-speech tasks, and suggest that the right hemisphere might be recruited to support sensorimotor processing in childhood stuttering. Differences in oscillatory dynamics occurred despite comparable task performance between groups, indicating that altered balance of cortical activity might be a core aspect of stuttering, observable during normal motor behavior.

## Introduction

Developmental stuttering is a neurodevelopmental disorder that affects the forward flow of speech, characterized by sound repetitions, prolongations, and blocks. Stuttering is a multifactorial disorder that arises from complex interactions between genetics, motor, language, cognitive, and emotion regulation abilities which impact neural development, especially in speech motor control networks (Smith & Weber, 2017). Stuttering affects ∼5% of preschool-age children and ∼1% of adults (Yairi and Seery, 2011) and is often associated with negative impacts on quality of life.

Stuttering has been associated with deficits in motor control, particularly for action initiation and timing (Alm et al., 2004; Civier et al., 2013; Chang and Guenther 2020). Besides stuttering events, people who stutter exhibit more variable speech movements compared to people who do not stutter even when their speech is perceptually fluent (e.g., Wiltshire et al., 2021; MacPherson & Smith., 2013; Smith et al., 2012). In non-speech motor tasks, people who stutter often perform slightly worse than people who do not stutter, particularly in demanding tasks that involve complex sequences of movements or fine timing control (e.g., Smith-Bandstra et al. 2006ab; Falk et al., 2014; Bauerly and De Nil, 2011; Toyomura et al., 2021), while results have been mixed for simple motor tasks, such as hand claps (Piispala et al., 2016; Hilger et al., 2016; Toyomura et al., 2021). These reported differences in speech and non-speech motor behavior could be related to subtle structural and functional differences in the sensorimotor system, including reduced size, activation and connectivity between the left prefrontal, premotor, motor and auditory cortices, as well as within the cortico-basal ganglia and cortico-cerebellar loops, and greater size, activation and connectivity in the right prefrontal and auditory cortices (Belyk et al., 2015; Brown et al., 2005; Chow and Chang 2017; Chang et al., 2019). Left hemispheric structural differences have been consistently reported both in children and adults who stutter, while right hemispheric differences have been typically found in adults. This suggests that the left hemisphere might be the core site of disruption in stuttering, while structural changes in the right hemisphere may develop as a compensatory mechanism over time, following repeated recruitment of these areas to support anomalous left hemisphere neural processes (e.g., Neef et al., 2011, 2015, 2018; Chang and Guenther, 2020). However, little is known about how the two hemispheres contribute to motor control in children who stutter, close to the onset of the disorder.

The neural mechanisms underlying motor control, including local neuronal processing and network properties, can be investigated non-invasively by measuring neural oscillations – endogenous periodic fluctuations in the excitability of neuronal populations – using electroencephalography, or EEG. (Buzsáki and Draguhn 2004; Lakatos et al., 2008; Sauseng and Klimesch, 2008; Fries 2005; Fries 2015). Neural oscillations in the beta range (13-30 Hz) have been particularly implicated in sensorimotor processing and motor control. Endogenous beta oscillations are present throughout the sensorimotor network, including cortical - somatosensory, motor and premotor - structures, as well as subcortical structures (Pfurtscheller and Da Silva, 1999; van Wijk et al., 2012). Cortical beta oscillations, which can be measured with the EEG, are reliably modulated during motor tasks: beta power is suppressed during movement planning and execution and enhanced after movement completion (Pfurtscheller and Da Silva, 1999). Reflecting the lateralization of the motor network, these changes in beta power are stronger in the left hemisphere during speech tasks, and in the hemisphere contralateral to the moving hand during unimanual tasks (Gehrig et al., 2012). According to one influential hypothesis, the suppression of beta power before and during motor activity may represent the release of motor inhibition necessary to initiate movement, while the increase in beta power after movement may actively maintain the existing motor state, inhibiting new movement initiation and facilitating the integration of sensory feedback (Engel and Fries, 2010; Baker, 2007). Research on motor disorders such as Parkinson’s disease support this interpretation, as difficulties to initiate movement has been associated with enhanced beta power at rest and reduced beta modulation during motor planning (e.g., Schnitzler and Gross, 2005; Heinrichs-Graham et al., 2014).

In stuttering research, beta power modulation has been investigated during speech planning and production. Although these studies are limited, they generally converge in reporting differences in beta modulation during fluent speech between adults who stutter and adults who do not stutter. However, the direction of the effect (increased or decreased beta modulation) and whether differences involved the left or right hemisphere, varied across studies (Rastatter et al., 1998; Salmelin et al., 2000; Sowman et al., 2012; Mersov et al., 2016, 2018; Jenson et al., 2018; Sengupta et al., 2017, 2019). These heterogeneous results might partly reflect the heterogeneity of the disorder across adult individuals who stutter, as well as differences in the level of difficulty of the speaking condition and in the linguistic computations that were required to perform the tasks across the different studies.

In addition to beta oscillations, slower oscillations in the delta (2-4 Hz) and theta (5-7 Hz) bands have been implicated in motor control, particularly in the coordination of the motor network. During movement, a distributed motor network becomes functionally integrated. The dynamic organization of the motor network is thought to be orchestrated by interregional synchronization of slow oscillations in the delta and theta bands (Babiloni et al., 2017; Rosjat et al., 2018, 2020; Yeom et al., 2020). Beta oscillations in different regions of the sensorimotor cortex have also been shown to transiently synchronize during motor tasks, although their role in the functional coupling of the motor network is not fully understood (Gross et al., 2005; Gerloff et al., 1998).

In contrast to the studies assessing beta power as a marker of speech motor processing, oscillatory connectivity during speech and non-speech motor tasks has not been investigated in stuttering. However, at rest, adults who stutter were reported to exhibit anomalous interhemispheric connectivity in multiple frequency bands compared to adults who do not stutter (Joos et al., 2014).

In sum, studies of oscillatory dynamics indicate atypical patterns of sensorimotor beta rhythm and altered resting state oscillatory connectivity in adults who stutter compared to adults who do not stutter. However, it is unclear whether differences in neural oscillations during speech or non-speech motor tasks are also present in children who stutter. To date, studies of oscillatory activity in children who stutter have been limited to altered beta oscillatory activity at rest or during in a rhythm listening task (Etchell et al., 2016; Ozge et al., 2004).

In the current study, we fill this critical gap in the literature by assessing whether spatiotemporal patterns of sensorimotor activity (as reflected in oscillatory power) and functional coupling (as reflected in oscillatory phase synchronization) around motor action initiation differ between children who stutter and children who do not stutter (ages 5 to10 y). To examine whether general differences in sensorimotor activities are present in stuttering (that are not limited to speech), we examined neural oscillations elicited by a non-speech motor task. Furthermore, to promote comparable performance across groups, we chose a simple task, consisting of pressing a button in response to an auditory target, which did not require complex movements or fine timing control. We measured temporal changes in beta power and in delta, theta and beta synchronization before and after the manual button press and compared these measures between left and right sensorimotor cortices and between children who stutter and children who do not stutter.

Based on previous results in adults who stutter and on theoretical accounts of stuttering that suggest left hemisphere as the core site of disruption, we predicted differences in power modulation and phase synchronization in the left hemisphere (contralateral to the response hand), but potentially not in the right (ipsilateral) hemisphere or between hemispheres.

Considering the simplicity of the task, we did not expect to see behavioral differences in average accuracy or response time. However, given previous results of heightened variability in the timing of movement initiation in both speech and non-speech motor tasks (e.g. Falk et al., 2014; Wiltshire et al., 2021), we expected to see greater variability in response time in the stuttering group. Greater neural variability, indicated by greater trial-by-trial variability in slow oscillatory dynamics (Popovych et al., 2016; Hamel-Thibault et al., 2018), may underlie response time variability. Thus, we predicted reduced consistency of phase locking to button press across trials in the delta and theta oscillations measured over the left hemisphere.

## Materials and Methods

### Participants

All study procedures were approved by the Human Research Protection Program at Michigan State University. Prior to data collection, all children’s primary caregivers gave written informed consent, and children gave assent to participate in the study.

Participants were 16 children with developmental stuttering (6 males) and 18 typically developing children (12 males), ages 5-10 years. All children were monolingual speakers of English, with no neurological deficit according to parental report. Children were included in the stuttering group based on the report of their parents and confirmation by a certified Speech-Language-Pathologist. Stuttering severity was measured by the Stuttering Severity Instrument, 4th Edition (SSI-4, Riley, 2009), with scores in the group of children who stutter ranging from very mild to severe (range: 6-28; median: 18; interquartile range 13-21). Three children were categorized as very mild (SSI-4 score ≤ 10), 7 as mild (SSI-4 score from 11 to 20), 5 as moderate (SSI-4 score from 21 to 27) and 1 as severe stuttering (SSI-4 score = 28).

All participants completed a battery of behavioral speech-language assessments (Table 1). These included measures of non-verbal intelligence (either the Primary Test of Non-verbal Intelligence, PTONI, Ehrler & McGhee, 2008, or the Test of Non-verbal Intelligence, TONI-4, Brown et al., 2010, based on age at time of testing), and receptive and expressive language (Clinical Evaluations of Language Fundamentals Preschool-2, second edition (CELF-P2, Wiig et al., 2004), or Clinical Evaluations of Language Fundamentals, Fifth Edition (CELF-5, Wiig et al., 2013) depending on age). All children scored within or above normal range, except one child who stutters who had lower performance on the PTONI (standard score: 64). Importantly, the two groups did not differ in these measures (nonverbal intelligence: *t*(32) = −1.08, *p* = .29; language: *t*(32) = .27, *p* = .79). All children completed a modified version of the Edinburgh Handedness Inventory test (Oldfield, 1971). All children were right-handed except for two ambidextrous children (one per group) and one left-handed child in the group of children who do not stutter. The groups did not differ in socioeconomic status (SES).^1^

**Table 1.** Demographics information for children in the two groups. No measure statistically differed between the groups. TONI = Test of non-verbal intelligence. PTONI = Primary test of non-verbal intelligence. CELF = Clinical Evaluation of Language Fundamentals, Fifth Edition, or Clinical Evaluation of Language Fundamentals-Preschool, Second Edition; SSI-4 = stuttering severity instrument 4. M=mean, SE=standard error.

|  | Children who stutter<br>M (SE) | Children who do not stutter<br>M (SE) |
| --- | --- | --- |
| Age (years) | 6.6 (.3) | 6.9 (.4) |
| Maternal education | 16.7 (.5) | 16.4 (.4) |
| Paternal education | 15.3 (.5) | 16.1 (.3) |
| Household income | 6.6 (.6) | 7.5 (.4) |
| TONI/PTONI | 116.2 (5.3) | 107.8 (5.6) |
| CELF | 113.19 (3.12) | 114.39 (3.26) |
| SSI-4 | 17.3 (1.6) | - |

### Task and stimuli

All children performed an auditory oddball task, in which they listened to a stream of frequent (standard, 1kHz) and rare (target, 2kHz) pure tones (ratio 80:20, Kaganovich et al., 2010). They responded to the rare tones by pressing a single button on a response pad with their right thumb (Current Design non-MR trainer response pad). Each tone was 50 ms long, including linear rise and fall time of 5 ms. Stimuli were presented via over-the-ear headphones at a comfortable loudness (∼75 dB SPL). Inter-stimulus intervals were 200 ms or 1000 ms, pseudorandomly presented such that rare target tones were always followed by the longer, 1000 ms interval to reduce overlap between motor responses and the subsequent tone stimulus. The task was administered in experimental blocks of 300 tones (240 standard and 60 targets per block). Children completed two blocks separated by a short break, except for 3 children in the non-stuttering group and 2 children in the stuttering group, who only completed one block. Brief rest breaks were available every 100 trials as needed. Experimental blocks were preceded by a short practice block (20 tones) to ensure children understood and were able to successfully complete the task. The practice block was repeated until a child could successfully performed the task (maximum practice blocks needed was 3). The groups did not differ in the number of practice or experimental blocks completed (experimental blocks, mean ± SE for both groups: 1.8 ± 0.1, *t*(32) = −0.08, *p* = 0.94; practice blocks, children who stutter: 0.9 ± 0.2, children who do not stutter: 1± 0.2, *t*(32) = 0.24, *p* = 0.81).

### Behavioral measures

For each participant, hit rate and false alarm rate were computed as the percentage of correct responses after a target tone (hit rate) and after a standard tone (false alarm rate). Response time was defined as the delay between the target onset and the button press on correct trials. For each participant, average response time and variability (standard deviation) of response times were computed. These measures of behavioral performance were compared across the two groups using t-tests.

### EEG recording and analyses

EEG data were continuously recorded at 512 Hz using 32 Ag/Ag-Cl electrodes embedded in an elastic cap (Biosemi Active-Two system), with locations consistent with the International 10-20 system. Additional electrodes were places over left and right mastoids, left and right outer canthi, and left orbital ridge.

EEG analyses were performed in using EEGLAB v2021.1 (Delorme and Makeig, 2004) and custom made routines in MATLAB R2016b (MathWorks, Natick, MA, USA). EEG recordings were referenced offline to the average of the left and right mastoids, filtered between 1 and 60 Hz and down sampled to 256 Hz. The portions of recordings collected during breaks were removed, leaving 2 sec buffer zones for edge artifacts. Remaining EEG data were inspected visually and with the aid of clean_rawdata EEGLAB plugin (https://sccn.ucsd.edu/wiki/EEGLAB_Extensions). Channels were removed if they had poor correlations with neighboring channels (below 0.85) or contained non-transient line noise greater than 4 standard deviations beyond the channel population mean. On average, 1.9 channels were removed per subject (most common: T7 and T8, over the temporal region). Time windows contaminated with artifacts were identified with artifact subspace reconstruction (ASR, Kothe and Jung, 2014), with a rejection threshold of 20. ASR has been shown to improve the quality of ICA decomposition (Chang et al., 2020; Anders et al., 2020). Oculomotor, muscle and channel artifacts were identified and subtracted from the data via Independent Component Analysis (ICA) in the EEGLAB toolbox. On average 2.3 components were removed after visual inspection of their topography, time course and power spectrum.

Only trials with correct motor responses were analyzed. For each subject, EEG data were segmented into 3 seconds epochs, from 1s before to 2s after the onset of correct button presses (epochs were 3s to allow space for edge artifacts induced by wavelet convolution). We analyzed the data within five standard frequency bands: delta (2-4 Hz), theta (5-7 Hz), alpha (8-12 Hz), beta (13-30 Hz) and gamma (32-50 Hz). Analyses were restricted to bilateral electrodes over the premotor and sensorimotor cortical areas (i.e., the frontocentral electrodes: F3, F4, FC5, FC6, C3, C4, C5, C6, Figure 1A) and to the time interval −200 to 200 ms around the button press, which comprises a preparatory period (−200 to 0 ms) and an execution period (0 to 200 ms). Due to the fast pace of the task, no stationary baseline (i.e., periods with no movement-related EEG signals) was present in the task, thus all analyses focused on the comparison between the execution and the preparatory periods.

**Figure 1.**
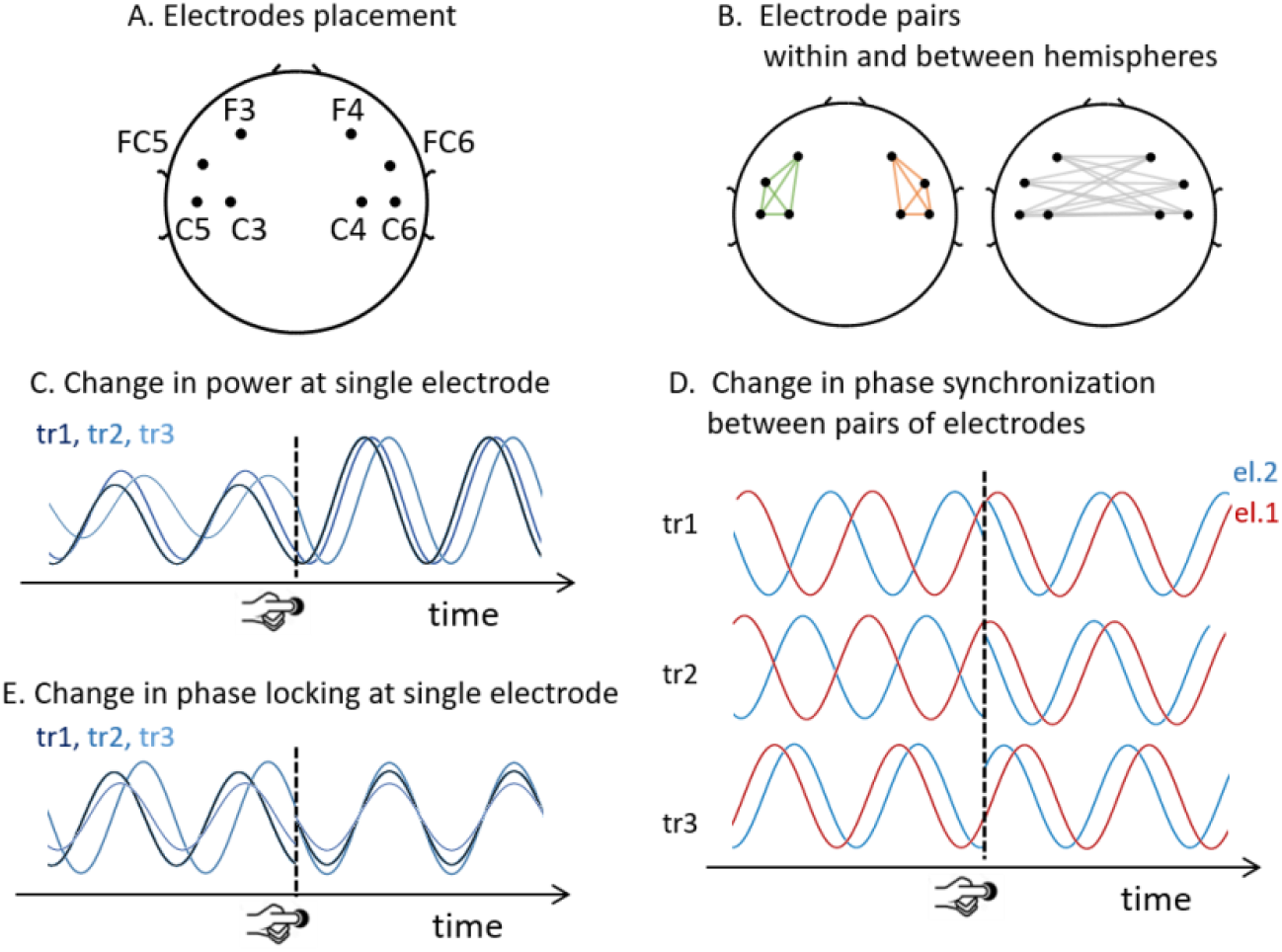
Electrode placement and measures of oscillatory activity. (**A**) All analyses included 8 bilateral frontocentral electrodes. (**B**) Within-hemisphere and between-hemisphere pairs of electrodes included in the analysis of phase synchronization. Green and orange colored lines connect each pair of electrodes within the left and right hemisphere, respectively (Left) and grey lines connect pairs of electrodes between hemispheres (Right). (**C**) Schematic of change in power with button press for three trials overlaid (tr1, tr2, tr3) (**D**) Schematic of change in inter-electrode phase synchronization for three trials stacked vertically (tr1, tr2, tr3). In each trial, the oscillations at two electrodes are indicated in red and blue (labelled el.1 and el.2). Across these trials, the phase difference between the two oscillatory signals is variable before button press and becomes more similar after button press, indicating an increase in phase synchronization. (**E**) Schematic of change in phase locking for three trials overlaid (tr1, tr2, tr3).

### Spectrotemporal analyses

Time-frequency decomposition was performed via complex Morlet wavelet convolution in the frequency range of 2-50 Hz (in 40 steps). The number of cycles in the wavelets increased linearly from 3 to 10. Figure 1 shows the three measures of oscillatory dynamics.

<u>Power</u> was computed from the wavelet transform at each frequency and time point. For each participant and for each electrode, power changes around button press were computed as the difference in decibels between the average power in the 200 ms after button press minus the average power in the 200 ms before button press (Figure 1C).

<u>Functional connectivity</u> between neural populations was measured as the <u>phase synchronization</u> between same-band oscillatory activity at two electrodes. Multiple indices have been proposed to measure phase synchronization (Bastos and Shoffelen, 2016). Here we chose the weighted phase-lag index (wPLI) because it is robust to volume conduction effects, that is, it does no yield false connectivity due to shared sources (Vinck et al., 2011, Cohen 2014; Bastos and Shoffelen, 2016). The wPLI measures the distribution of phase differences between two EEG signals across trials. To eliminate volume conduction effects, the wPLI disregards instantaneous coupling (i.e., it disregards phase angle differences equal to zero degrees) and weights phase angle differences farther from zero progressively more. The wPLI values range between 0 and 1, with 0 indicating no connectivity (i.e., a random distribution of phase differences over trials) and 1 indicating true lagged connectivity (i.e., the same non-zero phase difference in all trials). In particular, we measured the debiased weighted phase-lag index (dwPLI), which is an adjusted version of the wPLI index that is more robust to bias due to small sample size, as well as volume conduction and noise (Vinck et al., 2011, Cohen 2014; Bastos and Shoffelen, 2016). To evaluate changes in phase synchronization around the button press response, we computed the difference between the average dwPLI in the 200 ms after button press minus the average dwPLI in the 200 ms before button press for each frequency band (Figure 1B, D).

<u>Phase locking</u> was computed using the inter-trial phase clustering index (ITPC), which measures the consistency of an oscillation phase angle at a given time across trials (Cohen, 2014). ITPC values range between 0 and 1, with 0 indicating high variability of phases across trials, and 1 indicating no variability, i.e., all trials have the same phase. Changes in phase locking around the motor response were computed for each frequency band, as the difference between the average ITPC in the 200 ms after button press minus the average ITPC in the 200 ms before button press (Figure 1E).

### Statistical analysis

For each dependent variable (power, phase synchronization and phase locking) in different frequency bands, we performed a mixed-model ANOVA using a linear mixed-effects model (Murrar and Brauer, 2018). For the analysis of power and phase locking, fixed effects were Group (stuttering and non-stuttering) and Hemisphere (left and right). For the analysis of phase synchronization, fixed effects were Group (stuttering and non-stuttering) and Location (left hemisphere, right hemisphere, between-hemispheres). Subject was modeled as random intercept. Post-hoc comparisons were performed when the interaction and main effects were significant. Post-hoc comparisons were computed with F-tests on fixed-effects coefficients of the linear mixed-effects models using appropriate contrasts. p-values were adjusted for multiple comparisons with the Benjamini & Hochberg correction (Benjamini and Hochberg, 1995).

To investigate the relation between stuttering severity and oscillatory measures, we computed Pearson correlations between SSI-4 scores and average power, phase locking and phase synchronization in each hemisphere and between hemispheres for phase synchronization.

## Results

### 1. Behavioral performance was comparable in the two groups

Children who stutter and children who do not stutter exhibited comparable task performance in terms of hit rate, false alarms, average response time, and variability of the response time, as shown in Figure 2 and Table 2.

**Table 2.** Task performance for the two groups. M = mean, SE = standard error of the mean.

|  | Children who<br>stutter M (SE) | Children who<br>do not stutter<br>M (SE) | t-test |
| --- | --- | --- | --- |
| hit rate (%) | 72.86 (4.06) | 74.26 (2.41) | $t(32) = -.30; p = .76$ |
| false alarm rate (%) | 15.04 (3.72) | 7.65 (1.46) | $t(32) = 1.93; p = .06$ |
| average response time (ms) | 602.8 (22.7) | 628.8 (13.7) | $t(32) = 1.01; p = .32$ |
| variability of response time (ms) | 177.7 (11.4) | 153.7 (10.4) | $t(32) = -1.55; p = .07$ |

**Figure 2.**
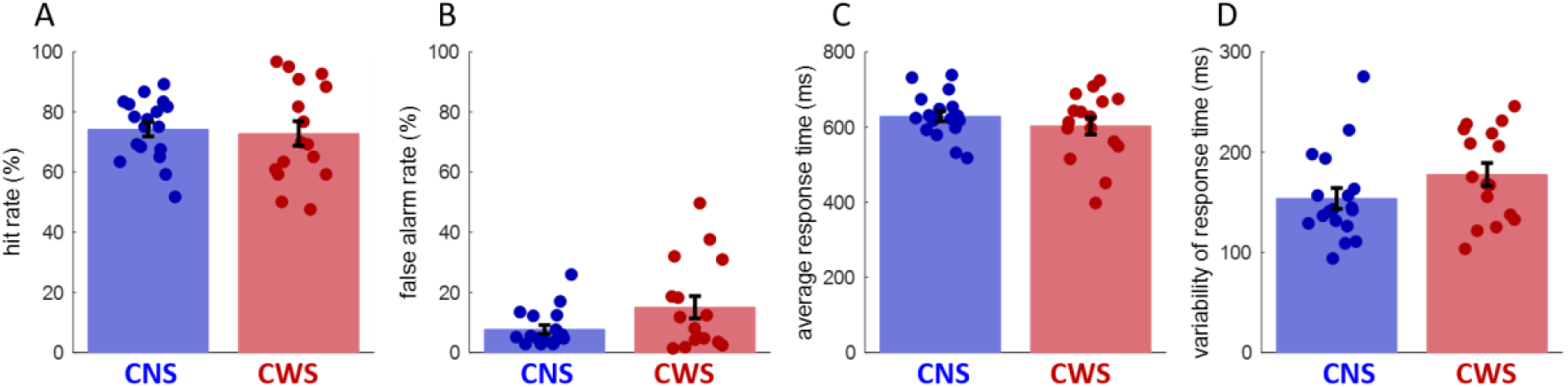
Comparable task performance in the two groups. (A) hit rate, (B) false alarms rate, (C) average response time, (D) variability of the response time (average standard deviation). Overlaid dots represent individual values. CNS = children who do not stutter, CWS = children who stutter.

### 2. Weaker beta power modulation over the left hemisphere in children who stutter

Figure 3A shows the average beta power over the left and right frontocentral regions (see EEG electrodes location in Figure 1) for the two groups. Children who do not stutter exhibited a brief beta power increase over the left hemisphere following button press response, which was not evident in children who stutter.

**Figure 3.**
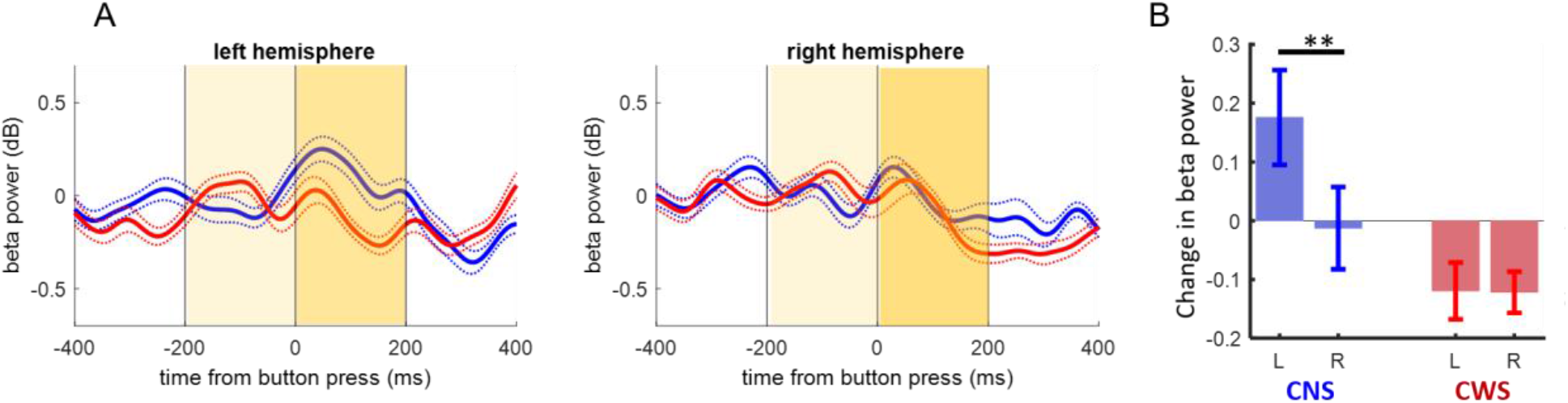
Movement related changes in beta power. (**A**) Time course of average beta power over the left and right hemispheres aligned to button press for children who do not stutter (CNS, blue) and children who stutter (CWS, red). Dotted lines indicate S.E.. Time windows for the analysis are indicated in yellow. (**B**) Changes in average beta power around button press (differences between average beta power in the 200 ms window after and the 200 ms window before the button press response). Beta power changes were left lateralized in children who do not stutter but not in children who stutter. Asterisks indicate significant results in post-hoc analyses (** *p* < 0.01).

We quantified beta power change around button press as the difference between the average beta power in the 200 ms window immediately after button press and the average beta power in the 200 ms window immediately before button press (time windows are indicated in yellow in Figure 3A). A mixed-model ANOVA revealed a significant interaction between group and hemisphere, indicating that power changes around button press were left lateralized in the non-stuttering group, but not in the stuttering group (*F*(1,268) = 4.34; *p* = .038, Figure 3B). In post-hoc analysis, the difference between the two groups in left hemisphere beta power changes did not survive adjustment for multiple comparisons (*F*(1,268) = 4.45, non-adjusted *p* = .036, adjusted *p* = .072), and the comparison of beta power changes between hemispheres revealed a left lateralization only in children who do not stutter (*F*(1,268) = 9.45; adjusted *p* = 0.009). We also saw a significant main effect of hemisphere (*F*(1,268) = 4.56; *p* = .034), which was driven by the non-stuttering group. The main effect of group was not significant (*F*(1,268) = 2.32; *p* = .13)

### 3. Oscillatory synchronization was weaker at low frequencies and less left lateralized in the beta band in children who stutter

To assess functional coupling within and between hemispheres, we analyzed phase synchronization in the delta, theta and beta bands. The time course of average phase synchronization over the left and right hemispheres and between hemispheres are shown in Figure 4A for each group. At low frequencies, in the delta and theta bands, phase synchronization increased soon after button press in the non-stuttering group more than in the stuttering group. In contrast, in the beta band, the magnitude of phase synchronization was overall comparable between the two groups, but the spatial pattern was different, with greater synchronization between hemispheres in the stuttering group and greater synchronization over the left hemisphere in the non-stuttering group.

**Figure 4:**
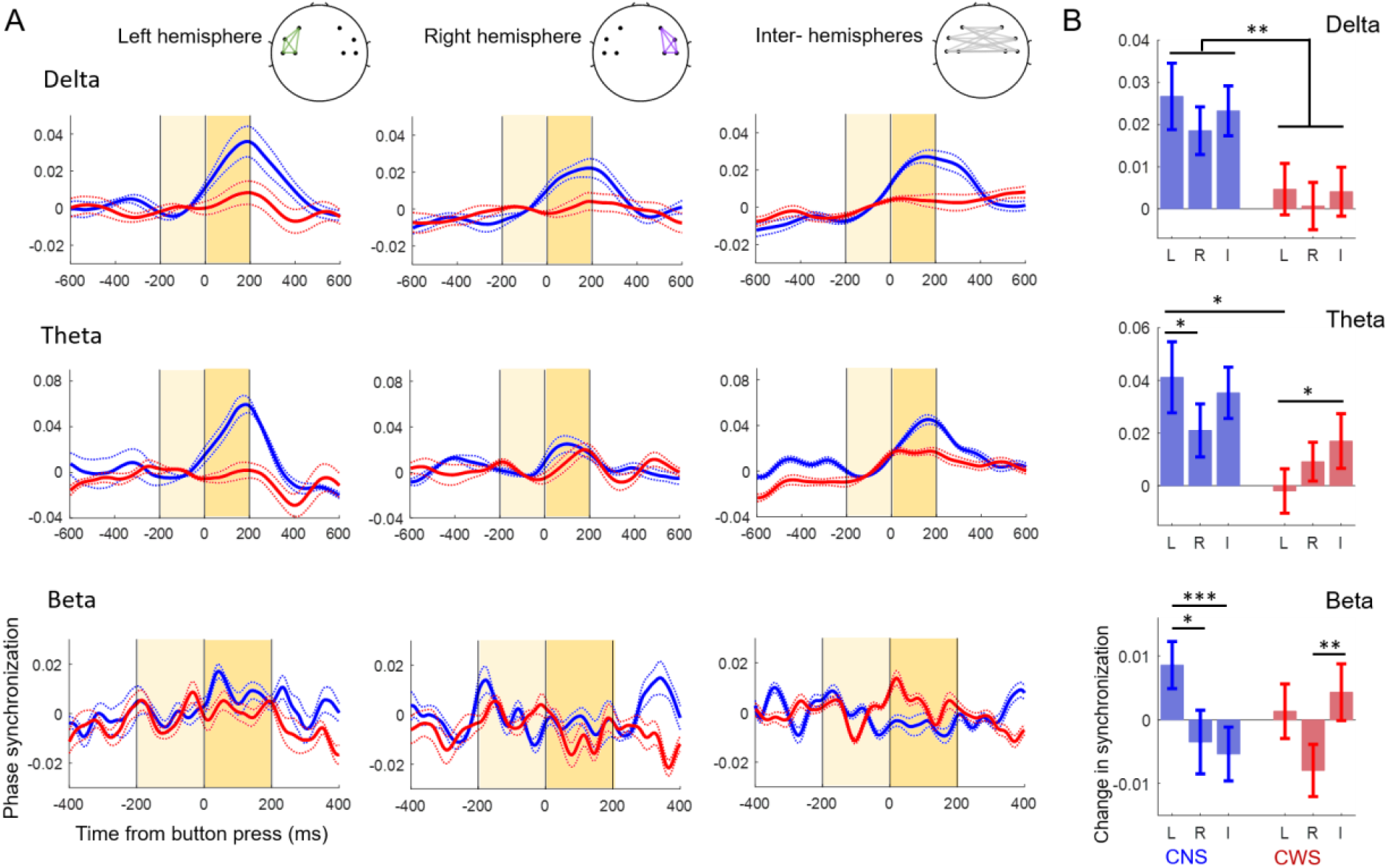
Changes in phase synchronization within and between hemispheres. (A) Time course of phase synchronization (dwPLI) within and between hemispheres in the delta, theta and beta bands for children who do not stutter (CNS, blue) and children who stutter (CWS, red). Dotted lines indicate S.E.. (B) In the delta and theta bands, changes in phase synchronization were greater in the children who do not stutter than in children who stutter. In the theta band, the difference between groups was greater over the left hemisphere. In the beta band, changes in phase synchronization were maximal over the left hemisphere in children who do not stutter, and between the two hemispheres in children who stutter. \**p* < .05, \*\**p* < .01, \*\*\**p* < .001, L = over left hemisphere, R = over right hemisphere, I = interhemispheric, or between, hemispheres.

Changes in phase synchronization around button press response (200 ms window after button press minus 200 ms window before button press) were analyzed with mixed-model ANOVAs in each band (Figure 4B, Table 3). In the delta and theta bands, we found a main effect of group, confirming that phase synchronization increased in the non-stuttering more than in the stuttering group (delta: *F*(1,946) = 9.84; *p* = .002, Theta: *F*(1,946) = 4.81; *p* = .029). In the theta band, we also observed a significant interaction between group and location (*F*(2,946) = 4.15; *p* = .016). Post-hoc comparisons showed that the difference between non-stuttering and stuttering groups was larger over the left hemisphere (*p* = .01; Table 4).

**Table 3.**
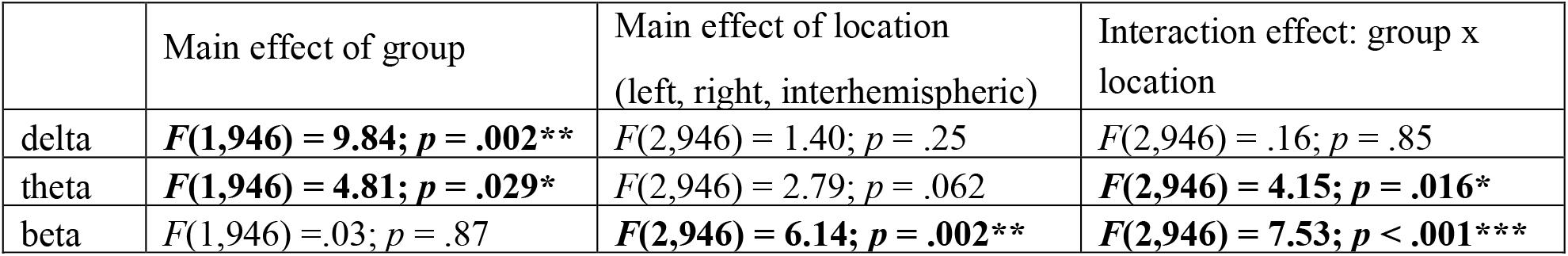
Results of linear mixed-model ANOVAs on phase synchronization to the button press response in the delta, theta and beta bands. Significant main effects and interactions are indicated in bold.

**Table 4:** Results of post-hoc comparisons of phase synchronization in the theta and beta bands using adjusted p-values (Benjamini & Hochberg correction). Significant comparisons are indicated in bold. L = over left hemisphere, R= over right hemisphere, I = interhemispheric, or between hemispheres.

|  | Between group comparison |  |  | Children who do not stutter |  |  | Children who stutter |  |  |
| --- | --- | --- | --- | --- | --- | --- | --- | --- | --- |
|  | L | R | I | L vs R | L vs I | R vs I | L vs R | L vs I | R vs I |
| Theta | **.01\*** | .38 | .20 | **.04\*** | .38 | .08 | .30 | **.04\*** | .35 |
| Beta | .28 | .47 | .06 | **.01\*** | **<.001\*\*\*** | .59 | .065 | .47 | **.004\*\*** |

In the beta band, we found a significant interaction between group and location, such that synchronization was left lateralized in the non-stuttering, but not in the stuttering group (*F*(2,946) = 7.53; *p* < .001). Post-hoc comparisons (Table 4) revealed stronger left hemisphere synchronization compared to synchronization over the right hemisphere or between hemispheres in children who do not stutter. In contrast, in children who stutter, left hemisphere and interhemispheric synchronization were comparable, and interhemispheric synchronization was stronger than right hemisphere synchronization. No significant differences in synchronization were observed between the two groups over left, right, or between hemispheres, but interhemispheric synchronization was marginally greater in children who stutter compared to children who do not stutter (Table 4).

### 4. Delta, but not theta, phase locking was less left lateralized in the children who stutter

To assess the cross-trial variability in oscillatory dynamics, we analyzed changes in slow oscillatory phase locking to button press. Figure 5A shows the time course of the average delta and theta phase locking in the left and right frontocentral regions. In both groups, phase locking to button press increases starting a few hundred milliseconds before the button press. Changes in phase locking (200 ms after minus 200 ms before button press) were analyzed with a mixed-model ANOVA in each band. In the delta band, there was a significant interaction between group and hemisphere (*F*(1,268) = 5.40; *p* = .021), indicating that the change in delta phase locking was left lateralized only in children who do not stutter. A significant effect of hemisphere (*F*(1,268) = 8.85; *p* = .003) was also observed, driven by the hemisphere differences in the children who do not stutter, as post-hoc analysis revealed left-lateralized delta phase locking only in the non-stuttering group (adjusted p-value of 0.006). In the theta band, there was a main effect of hemisphere (*F*(1,268) = 9.08; *p* = .0028) indicating that theta phase locking was left-lateralized in both groups (Figure 5B).

**Figure 5.**
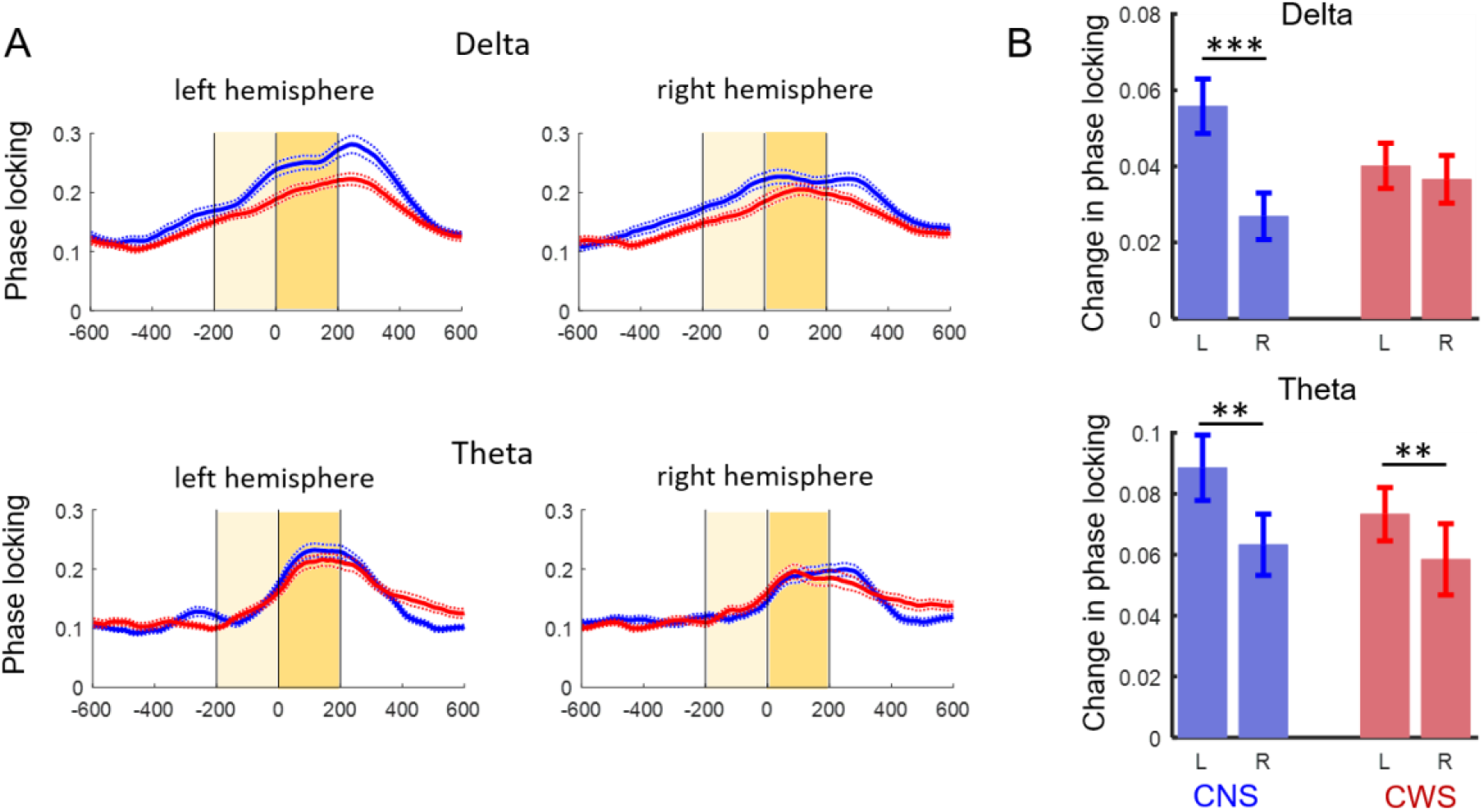
Phase locking to movement initiation. (**A**) Time course of average phase locking (ITPC) over the left and right hemispheres in the delta and theta bands for children who do not stutter (CNS, blue) and children who stutter (CWS, red). Dotted lines indicate S.E.. (**B**) Average changes in phase locking around button press response. In the delta band, phase locking was left-lateralized only in the non-stuttering group. In the theta band, phase locking was left lateralized in both groups. \*\**p* < .01, \*\*\**p* < .001, L = over left hemisphere, R = over right hemisphere.

### 5. Correlation between stuttering severity and measures of oscillatory dynamics

No correlation between stuttering severity (as measured by SSI-4 scores) and oscillatory measures survived multiple comparison, but uncorrected correlation analysis suggested a negative relation between stuttering severity and the average delta phase synchronization over the right hemisphere, such that children with more severe stuttering tended to exhibit reduced phase synchronization in the delta band over the right hemisphere (*r* = –0.53, *non-adjusted p* = 0.03, *adjusted p* > 0.05).

## Discussion

We investigated the spatiotemporal dynamics of sensorimotor oscillatory activity and connectivity around movement initiation in children who stutter and children who do not stutter. Children pressed a button in response to rare auditory targets embedded in a stream of non-target tones. Although task performance in terms of speed and accuracy was comparable between the two groups, the spatiotemporal dynamics of oscillatory activity and network connectivity during the non-speech movement differed. This suggests that differences in general sensorimotor control (non-speech) are present in children who stutter, and that the examination of oscillatory dynamics is a sensitive tool to investigate them.

### Behavioral performance

Given the simplicity of the task and required motor movement (responses consisted of a simple one-finger button press), we expected to find comparable behavioral accuracy and response speed, but we speculated that children who stutter might be more variable in response timing across trials. Indeed, consistent with our expectation and with previous studies that employed simple motor tasks (e.g., Piispala et al., 2016; Hilger et al., 2016), we found comparable accuracy and mean response time between the two groups. Children who stutter tended to be slightly more variable in their response time, but the difference did not reach significance (*p* = .07). This is consistent with previous reports of higher variability in the timing of movement initiation in both speech and non-speech motor tasks, which may result in significant group differences when performing more challenging tasks (e.g., Falk et al., 2014; Wiltshire et al., 2021; Smith et al., 2012).

### Beta power modulation

In this study, we found a lack of left lateralized peri-motor beta power modulation in children who stutter. After button press, beta power increased over the left hemisphere for children who do not stutter, while it slightly decreased over both hemispheres in children who stutter. This result is in general agreement with previous studies of oscillatory activity in adults who stutter, which have implicated altered modulation and lateralization of beta oscillations in the preparation and execution of fluent speech (Rastatter et al., 1998; Salmelin et al., 2000; Sowman et al., 2012; Mersov et al., 2016, 2018; Jenson et al., 2018; Sengupta et al., 2017, 2019). However, previous studies reported variation in the direction of the effect, with both stronger and weaker modulation of beta power being reported (e.g., Mersov et al., 2016; Jenson et al., 2018). In part, the inconsistencies in the results of previous studies with adults who stutter may reflect the heterogeneity of the disorder across individuals, including compensatory mechanisms, which may develop in adults who stutter either spontaneously or through therapy. In contrast, we have investigated children between 5 and 10 years of age, thus closer to the onset of the disorder.

Our results of weaker modulation of left peri-motor beta in the stuttering group are similar to results by Jenson and colleagues (2018) for speech production in adults who stutter, although methodological differences, including the nature and timing of the tasks (slow speech production vs. fast finger movement, adults vs. children) preclude a direct comparison. The reduced beta modulation found by Jensen and colleagues (2018) consisted of reduced beta suppression before and during speech movements over the left hemisphere, which was interpreted as reflecting reduced activation of the forward model of speech; however, post-movement beta rebound was not evaluated. In contrast, in our investigation, the reduced beta modulation in children who stutter was driven by a weaker rebound of beta activity after button press (see the time course of the beta power in Figure 3), although we cannot fully exclude that a difference was also present before movement (given the lack of a resting condition, it was not possible to independently estimate the levels of pre-movement suppression and subsequent rebound of beta activity). Indeed, studies of repetitive movements have shown that beta modulation comprises both a sustained suppression lasting the entire movement period, as well as amplitude modulations around each individual movement (e.g., Seeber et al., 2016).

The role of post-movement beta rebound is not fully understood, but studies suggest that its modulation is to some extent independent of pre-movement beta decreases (e.g., Hall et al., 2011). According to two competing, but not mutually exclusive hypotheses, beta rebound may reflect the active inhibition of the motor cortex to suppress further movement (Salmelin et al., 1995; Pfurtscheller et al., 1996; Schmidt et al., 2019) and/or it may reflect the post-movement processing of sensory feedback to evaluate movement outcome (e.g., Saltuklaroglu et al., 2018). Furthermore, an EEG study that manipulated task and sensory uncertainty found that the amplitude of beta rebound decreased with increasing uncertainty in feedforward predictions (Tan et al., 2016). Thus, reduced beta rebound is consistent with theories on stuttering that suggest unstable internal models and/or higher reliance on sensory feedback in people who stutter compared to people who do not stutter (e.g., Max et al., 2004; Civier et al., 2010). Future studies are necessary to distinguish between these hypotheses.

### Oscillatory synchronization

The second main difference in oscillatory dynamics concerns peri-movement changes in oscillatory phase synchronization, a measure of dynamic functional organization of the sensorimotor network (Fries 2005; Engel and Singer 2001; Varela et al. 2001; van Wijk et al., 2012). Consistent with previous studies in unimanual motor tasks (Ohara et al. 2001; Ramayya et al., 2020; Rosjat et al., 2018), typically developing children displayed widespread increase in slow frequency synchronization (delta/theta) and a left-lateralized pattern of beta synchronization. In the stuttering group, we expected to see less phase synchronization increase over the left hemisphere compared to the non-stuttering group, reflecting reduced functional connectivity within the left sensorimotor network during movement planning and execution. We found a complex pattern of results that partially agreed with our predictions. Relative to children who do not stutter, children who stutter displayed smaller peri-movement increase in connectivity at slow frequencies (delta/theta), but this difference was widespread, over left and right hemisphere as well as between hemispheres. In the beta band, children who stutter displayed less left lateralized connectivity than children who do not stutter, but the difference was driven by an increase in interhemispheric synchronization rather than by a decrease in left hemisphere synchronization.

Phase synchronization in delta/theta bands has been linked to long distance communication in the bilateral sensorimotor network (e.g., Rosjat et al., 2018; van Wijk et al., 2012). Network coupling has been proposed to rely on the interaction of two oscillatory mechanisms: long-range, low frequency phase synchronization and local hierarchical coupling between slow oscillations and fast oscillations/neural spiking. Within the cortical striatal network, invasive studies in animal models (e.g., Igarashi et al., 2013; von Nicolai et al., 2014; Zeng et al., 2021) support this proposed mechanism, particularly involving theta phase synchronization and theta-gamma coupling, while evidence for delta phase synchronization and delta-beta/gamma coupling is sparser (Dejean et al., 2011; von Nicolai et al., 2014; Morillon et al., 2019). Human non-invasive studies have related changes in delta/theta interregional synchronization with different phases of movements, such as preparation and execution (Rosjat et al., 2018, 2020; Yeom et al., 2020), and have highlighted the role of theta-gamma and theta-beta coupling in sensory feedback processing during speech production (Sengupta and Nasir, 2015; Sengupta et al., 2019). In addition, a computational model of the cortical striatal network has proposed that theta oscillations (with nested gamma oscillations) may encode motor sequences, similar to the encoding of spatial information in the hippocampus (Fukai et al., 1999). Thus, reduced delta and theta synchronization during movement in children who stutter may indicate less efficient communication in the motor network, which could particularly affect motor planning in complex sequences, such as speech.

The smaller increase in global delta/theta synchronization was accompanied by a larger interhemispheric beta synchronization increase in children who stutter. Interhemispheric beta synchronization has been shown to transiently change with movement initiation, increasing during coordinated bimanual tasks and decreasing during unimanual tasks (Serrien and Brown 2002; Ohara et al. 2001, de Oliveira et al., 2001; van Wijk et al. 2012). Our results suggest the possible recruitment of the right hemisphere for children who stutter during a unimanual simple task performance. However, more studies are necessary to understand how fast and slow oscillatory connectivity contribute to motor control in typically developing children and in children who stutter.

To our knowledge, no other study has examined changes in within-frequency, long range phase synchronization during motor tasks in stuttering. At rest, high frequency interhemispheric synchronization has been reported to be smaller in adults who stutter than in adults who do not stutter, and within the group of stuttering adults, more severe stuttering was associated with stronger interhemispheric low frequency synchronization (Joos et al., 2014). Methodological differences (adults vs. children and resting-state vs. task-evoked connectivity) preclude a direct comparison between our results and those by Joos et al., 2014; however, both studies point to altered network organization in stuttering that merits further investigation.

### Low frequency phase locking

We predicted that children who stutter would exhibit greater trial-by-trial variability in slow oscillatory dynamics over the left hemisphere, as indexed by a weaker transient phase locking to button press in the delta and theta bands. Results in the delta band but not in the theta band followed our prediction. In children who do not stutter, transient increase in delta and theta phase locking to the impending button press was stronger over the left hemisphere (contralateral to the moving hand). These patterns are consistent with previous studies of finger movements in children and adults (Gaetz et al., 2010; Popovych et al., 2016; Armstrong et al., 2018). In children who stutter, phase locking transiently increased with the same dynamics in both delta and theta bands, but in the delta band, it was not left lateralized; increases were similar across both hemipsheres. Although, as already noted, children who stutter were only marginally more variable in their response times than children who do not stutter, the observed weaker phase locking over the left hemisphere may reflect greater variability in the timing of motor activity.

## Limitations

This study has several limitations. The first is the number of participants. Developmental stuttering is a disorder with a complex etiology that involves the interaction of multiple factors, hence even at the onset of the disorder, children who stutter might differ in the neural processing supporting motor behavior. A larger number of participants is necessary to replicate and expand the results reported here. A second limitation of this study is that we did not collect data on the button press movement itself. We cannot rule out that different oscillatory dynamics between groups might have resulted from differences in movement parameters, such as velocity, duration of the press and force applied to the button, which we did not measure, as we focused on changes in oscillatory measures around movement onset. However, it is possible that these aspects of movement may contribute to the dynamics of oscillatory activity in complex ways. Finally, it is important to note that the oscillatory activity in our task might reflect sensorimotor as well as other cognitive processing, such as memory, attention and effort (van Wijk et al. 2012). Although these cognitive operations cannot be fully disambiguated, our results suggest that children who stutter and typically developing children differ in oscillatory dynamics around movement initiation, and in particular in the balance between the different frequency bands and the two hemispheres. Future studies involving diverse tasks and multiple methodologies may clarify the contribution of cognitive processes to these differences.

## Conclusions

The onset of movement is reliably accompanied by changes in oscillatory activity in the motor cortex, particularly in the hemisphere contralateral to the moving hand. In this study, we identified several differences in oscillatory dynamics between children who stutter and children who do not stutter, indexed by reduced modulation of power and phase synchronization over the left hemisphere, reduced lateralization of phase locking to movement initiation at slow frequencies, and altered interhemispheric phase synchronization in the delta and beta bands. Taken together, these results point to deficits in left sensorimotor cortex function in stuttering that may not be limited to speech, and suggest that the right hemisphere may be recruited to support sensorimotor processing even in childhood developmental stuttering.

## Footnotes

1 SES was measured by assessing maternal and paternal education as well as household income in the previous year. Maternal and paternal education was coded on a 1 to 18 scale based on years of education as follows: 10 = some high school; 12 = completed high school; 13 = partial college; 14 = 2-year college degree; 16 = standard college/bachelor degree; and 18 = graduate school. Household income was coded on a 1 to 11 scale based on the total income in the previous year as follows: 1 = less than 10,000 $; 2 = 10,000 to 25,000 $; 3 = 25,000 to 40,000 $; 4 = 40,000 to 55,000 $; 5 = 55,000 to 70,000 $; 6 = 70,000 to 85,000 $; 7 = 85,000 to 95,000 $; 8 = 95,000 to 105,000 $; 9 = 105,000 to 150,000 $; 10 = 150,000 to 250,000 $; 11 = more than 250,000 $ (e.g., Pollak and Wolfe, 2020).

